# Interaction of spatial attention and the associated reward value of audiovisual stimuli

**DOI:** 10.1101/2023.06.18.545478

**Authors:** Roman Vakhrushev, Arezoo Pooresmaeili

## Abstract

Reward value and selective attention both enhance the representation of sensory stimuli at the earliest stages of processing. It is still debated whether and how reward-driven and attentional mechanisms interact to influence perception. Here we ask whether the interaction between reward value and selective attention depends on the sensory modality through which the reward information is conveyed. Human participants first learned the reward value of uni-modal visual and auditory stimuli during a conditioning phase. Subsequently, they performed a target detection task on bimodal stimuli containing a previously rewarded stimulus in one, both, or neither of the modalities. Additionally, participants were required to focus their attention on one side and only report targets on the attended side. Our results showed a strong modulation of visual and auditory event-related potentials (ERPs) by spatial attention. We found no main effect of reward value but importantly an interaction effect was found as the strength of attentional modulation of the ERPs was significantly affected by the reward value. When reward effects were inspected separately with respect to each modality, auditory value-driven modulation of attention was found to dominate the ERP effects whereas visual reward value on its own led to no effect, likely due to its interference with the target processing. These results inspire a two-stage model where first the salience of a high reward stimulus is enhanced on a local priority map specific to each sensory modality, and at a second stage reward value and top-down attentional mechanisms are integrated across sensory modalities to affect perception.

## Introduction

Our surrounding environment contains a large amount of information whereas our brain’s processing capacity is limited. An important mechanism to face this challenge is to prioritize the processing of information that is relevant to our current goals and is associated with the most valuable outcome. However, these aspects, i.e., relevance and reward value, may not be always in the same direction. For instance, when waiting for our friend to pick us up at an intersection, we try to focus our attention on the side where our friend is most likely to arrive. At the same time, we may also notice the sound or sight of approaching vehicles that deliver takeout food, and if hungry we may be even more sensitive to these sources of information compared to the familiar sight of our friend’s car. Due to the prevalence of such situations, where environmental stimuli should be processed both based on their relevance as well as their reward value, a large body of literature has sought to investigate the underlying mechanisms of selective attention and reward-driven modulation of perception, as well as the interaction between the two (Maunsell, 2004; Carrasco, 2011; Chelazzi et al., 2013; Pessoa, 2015; Anderson, 2016a; Failing and Theeuwes, 2018; Anderson et al., 2021). However, the exact nature of this interaction and its underlying mechanisms have remained unknown.

Converging evidence from behavioral and neurophysiological studies has shown that reward value modulates sensory perception and its neuronal correlates (Serences, 2008; Hickey et al., 2010; Baldassi and Simoncini, 2011; Hughes et al., 2013; Leo and Noppeney, 2014; Pooresmaeili et al., 2014; San Martín et al., 2016; Bayer et al., 2017). Similarly, selective attention affects behavioral and neural responses to sensory inputs (Desimone and Duncan, 1995). The similarity between reward and attention has raised the question of whether they can be dissociated from each other at all (Maunsell, 2004). Previous studies have provided divergent answers to this question. On the one hand, the majority of past studies have reported a strong interaction between reward and attention (Yantis et al., 2012; Anderson, 2013; Chelazzi et al., 2013; Le Pelley et al., 2016; Failing and Theeuwes, 2018), suggesting a *dependence* between the two. For instance, neurophysiological studies showed similar effects of reward and attention on neuronal responses of area V1, and that reward effects are gated by attention, inspiring the idea of a ‘unified selection signal’ comprising both factors (Stanisor et al., 2013). Other studies showed a different form of interaction where the strength of attention to locations or features of stimuli was gated by reward (Chelazzi et al., 2013, 2014; Pessoa, 2015). These studies hence point to a bi-directional interaction between attention and reward, each influencing the strength of the other. This view is also in line with recent suggestions that reward and attention jointly influence stimulus representations on *an integrated priority map* (Chelazzi et al., 2014; Failing and Theeuwes, 2018), although it is unknown how such an integration is implemented. In contrast to this idea, yet another set of studies provided evidence for the *independence* of reward and attention, as reward effects were unchanged by the amount of attentional load (Baldassi and Simoncini, 2011) or the relevance of reward cue to the task (Garcia-Lazaro et al., 2018) occurring even rewarded stimuli were outside of conscious awareness (Lunghi and Pooresmaeili, 2023), and were observed during separable stages of neural processing (Bayer et al., 2017).

Similar to the approach of the latter studies, one possible way to dissociate the effects of reward and attention is to manipulate each factor orthogonally to the other. For instance, if successful detection of a target among distractors leads to the delivery of a reward, target features that are predictive of higher reward not only signal a better outcome (i.e., reward) but also engage attentional mechanisms more strongly than low reward stimuli. However, incidental reward stimuli that are linked to past rewarding experiences or predict reward magnitude independent of the performance in a task are more likely to have less dependence on task-related attentional processes. Likewise, orthogonal manipulation of reward and attention can be achieved when reward information is signalled through a stimulus feature (Baldassi and Simoncini, 2011; Garcia-Lazaro et al., 2018) or sensory modality (Pooresmaeili et al., 2014; Vakhrushev et al., 2021; Antono et al., 2022) that is distinct from the task-relevant target. Combining these, i.e., using previously rewarded stimuli that are task-irrelevant and are delivered through a different sensory modality, is likely to allow the maximum separation between the reward-related and the attentional prioritization of stimulus processing. In the current study, we try to shed light on the interaction between reward and attention under such conditions.

We draw on previous findings showing that reward stimuli from a different sensory modality (audition) can affect perception in vision (Pooresmaeili et al., 2014; Antono et al., 2022), even when cross-modal reward stimuli (auditory sounds) are irrelevant to the task at hand (i.e., visual orientation discrimination). Since there is evidence for the separation of attentional resources across sensory modalities (Duncan et al., 1997; Alais et al., 2006), these findings may indicate that cross-modal reward stimuli affect visual processing independently of visual attention. More recently (Vakhrushev et al., 2021), we showed that behavioral and EEG correlates of reward effects differ between stimuli that are in the *same* or *different* sensory modalities as the target (intra-modal and cross-modal, respectively). Whereas intra-modal reward stimuli led to an early suppression of the visual event-related potentials (ERPs), cross-modal rewards boosted the visual ERPs later in time and more persistently. These results further support the idea that the interaction of attentional and reward mechanisms may depend on the sensory modality of the reward with respect to the target of the task. However, in the latter study, intra-modal and cross-modal conditions not only differed in how reward was cued but also in whether they involved the processing of a unimodal (only visual) or a bimodal (audiovisual) stimulus. Additionally, in these previous studies selective attention was not systematically manipulated, and therefore the assumed interaction between the sensory modality and attention could not be directly tested.

In the current study, we remedy the shortcomings mentioned above and ask whether the behavioral and neural effects of reward and attention occur independently from each other. Additionally, we ask whether a putative interaction between attention and reward depends on the sensory modality of reward stimuli. To answer these questions, we modified a behavioral paradigm developed by a previous study Talsma (Talsma and Woldorff, 2005) where it is possible to independently control both attention and reward. In this paradigm, participants performed a target detection task on bimodal (audiovisual) stimuli. Spatial attention was manipulated in a block-wise manner, where in each block participants were asked to attend either to the left or to the right visual hemifield and report changes that occurred in the attended hemifield. Reward value was manipulated by associating a specific feature of visual (tilt orientation) or auditory (pitch) stimuli with different magnitudes of monetary reward (high-value or no-value), resulting in stimuli where either one, both or neither of the modalities were associated with reward. Importantly, these features were orthogonal to the target detection task and were not predictive of the reward delivery as reward associations were learned in a separate conditioning task prior to the test phase. Concurrently with this task, EEG data was recorded allowing us to inspect the effects of reward and attention on visual as well as auditory ERPs. All procedures and hypotheses (referred to as H1-H7) were preregistered (https://osf.io/xte4v).

We expected to find that both reward value (H1) and allocation of attention (H2) enhance the behavioral and electrophysiological indices of sensory perception. Additionally, we predicted that there would be no significant interaction between reward and attention (H3). The latter hypothesis was based on two lines of reasoning: firstly, by maximal orthogonalization of reward and attention we predicted that both factors could independently affect sensory perception. Secondly, the task imposed a minimum load on the attentional control as each trial only contained one stimulus and attention was manipulated across blocks. These characteristics were optimal for allowing the reward information to influence perception both on the attended as well as on the unattended side without a significant cost for the system, hence leading to reward modulations independently of attention. Additionally, we hypothesized that the reward and attention effects and their interaction might depend on the sensory configuration of the reward stimuli, i.e., whether visual, auditory or both visual and auditory stimuli, are associated with reward (H4-H6). Specifically, stimuli containing high reward stimuli in both modalities were predicted to produce the strongest value-driven effects compared to those that contained unrewarded stimuli. Additionally, we expected similar reward-driven and attentional effects on the ERPs originating from the visual or auditory cortex (H7). To provide a preview of our results, we found no main effect of reward value (rejecting H1), a strong effect of attention (confirming H2), and an interaction between reward and attention (rejecting H3) where attentional modulation of ERPs was strongest for the high reward stimuli. Additionally, the latter effect showed a dependence on the sensory modality, with the strongest effects observed for the stimuli which contained an auditory high reward stimulus (confirming H6) across all brain areas (confirming H7).

## Methods

### Participants

The sample size was based on a previous study (Vakhrushev et al., 2021) and was set to N = 36, but to compensate for possible dropouts and outliers we recorded data from N = 42 participants. Four participants were removed from the analysis due to their low performance during the behavioral task (accuracy < 60%) resulting in the data from 38 participants to be included in our analyses (age: 27.2 ± 5.4 years -mean ± SD-; 19 females; 4 left-handed). Subjects were recruited via a local database and had normal or corrected to normal vision. Before the experiment started and after all procedures were explained, participants gave an informed written consent and participated in a practice session. All procedures were approved by “Universitätsmedizin Göttingen” (UMG) under proposal number 15/7/15.

### Stimuli and task

The behavioral paradigm employed during the conditioning (**Figure 1a**) and the main task (**Figure 1b**) were adopted from a previous study (Talsma and Woldorff, 2005) and involved the detection of infrequent targets in a rapidly presented stream of unimodal (either auditory or visual during the conditioning) or bimodal (audiovisual during the main task) stimuli. All stimuli were lateralized 15° relative to the vertical meridian and 6° below the horizontal meridian, each presented briefly for a duration of 108.3 ms. Auditory stimuli were either a 1050 Hz or a 350 Hz tone with a linear rise and fall of 10 msec and an amplitude of 75 dB which were convolved with head-related transfer function (HRTF) filters to render them co-localized with the visual stimuli (Algazi et al., 2001) and were delivered through in-ear headphones. Visual stimuli were tilted square-wave gratings (6° visual degrees or about 5.8 × 5.8 cm), oriented ±45° and modulated between white and gray colors presented on a grey background. Audiovisual stimuli contained auditory and visual stimuli presented synchronously and at the same location producing the impression of one single bimodal object.

**Figure 1.**
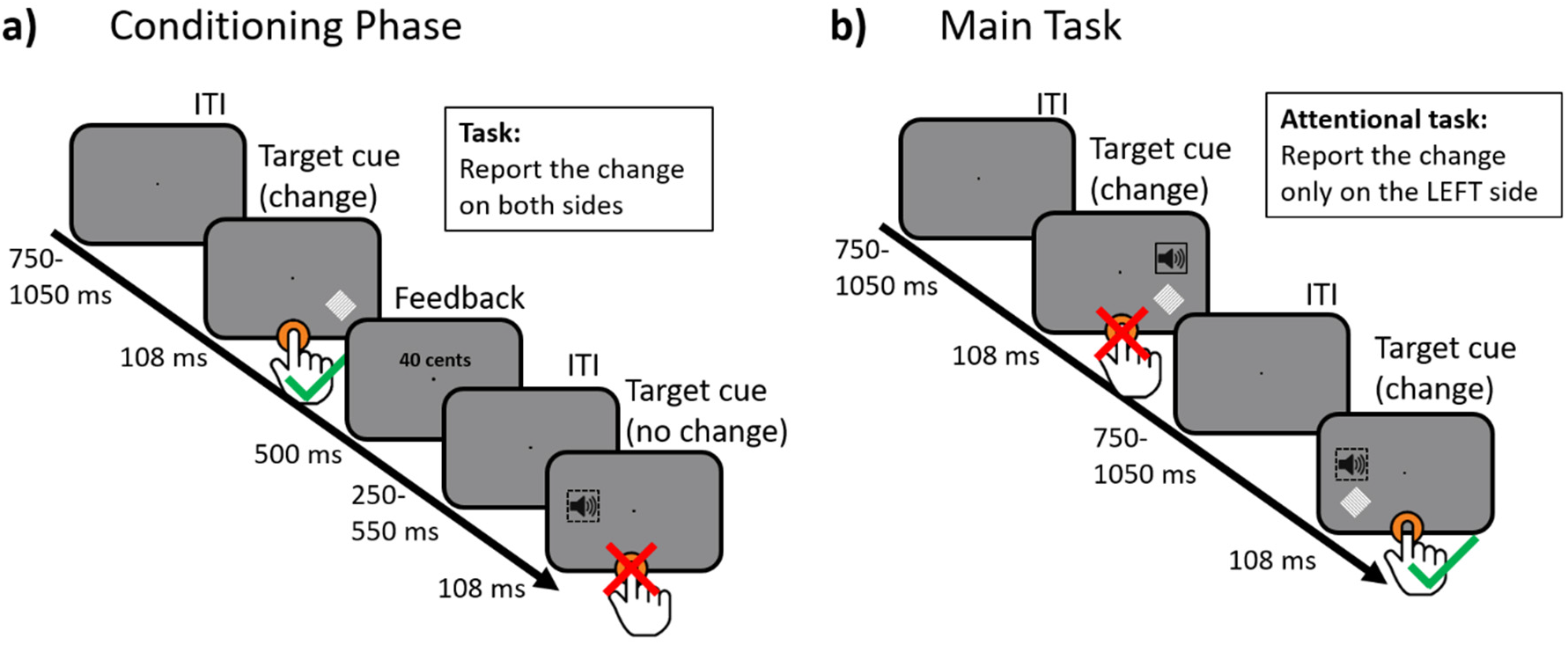
Behavioral paradigm. Participants were instructed to detect occasional targets by pressing a “space bar button”. Targets were trials with a change in intensity of auditory and/or visual stimuli midway through their presentation. The change made an impression of a stutter (auditory condition) or a flicker (visual condition). Both visual (±45° tilt) and auditory stimuli (350Hz and 1050Hz) were presented peripherally (15°) and below the fixation point (6°). **a)** The experiment started with a conditioning phase in which reward associations were learned. **b)** In the main task, visual and auditory stimuli always appeared together: unilaterally and synchronously. At the beginning of every block with the main task, participants were instructed to pay attention only to one side of the screen and ignore target trials on the other side (in the demonstration above participants were instructed to only report changes on the left side).

Throughout the experiment, the participants’ task was to detect target trials, i.e., trials containing a change, by pressing a “Space” button on a keyboard. Target trials were randomly interleaved with the rest of the trials and constituted 20% of all trials. Target trials were similar to standard trials, except that halfway through the presentation of stimuli a transient drop of stimulus intensity occurred, which caused an impression of the stimulus having a stutter (auditory stimuli) or a flicker (visual stimuli).

### Conditioning phase

All trials in the conditioning phase contained unimodal (only visual or only auditory) stimuli and were grouped into blocks of 27 trials by the modality. Blocks with different modalities were pseudo-randomly interleaved across the conditioning phase. Participants were instructed to report target trials (trials with a transient change) on both sides of the screen.

During the conditioning phase, participants learned to associate different tones and line orientations with either a positive or zero monetary reward (high-value or no-value, respectively). To achieve this goal, all trials in conditioning phase were followed by a feedback display (500 ms) showing the monetary reward outcome assigned to the presented stimulus. These rewards were drawn from two Poisson distributions with the mean of 40 cents and SD of 5.5 cents for high-reward trials (maximum reward was fixed at 50) and 0 cents for no-reward trials. Participants were instructed to remember the reward outcome of each condition and later report it (at the end of each block in the conditioning phase, through a 2AFC procedure in which the high reward stimulus of two consecutively presented stimuli had to be reported).

In total, the conditioning phase had *four* conditions: (1) high-value visual, (2) High-value auditory, (3) no-value visual, and (4) no-value auditory (**Figure 2a**). These conditions were divided into factors *modality* (visual or auditory) and *reward* (high- or no-value). Participants completed 960 trials during the conditioning phase, where each condition was repeated 240 times (120 trials per screen side). 20% of all trials were targets (trials with a transient change). Finally, to control for the perceptual biases towards visual and auditory stimuli, this phase was repeated twice and in the second repetition the reward pairings were reversed. This was indicated to participants with an instruction display before the second repetition commenced.

**Figure 2.**
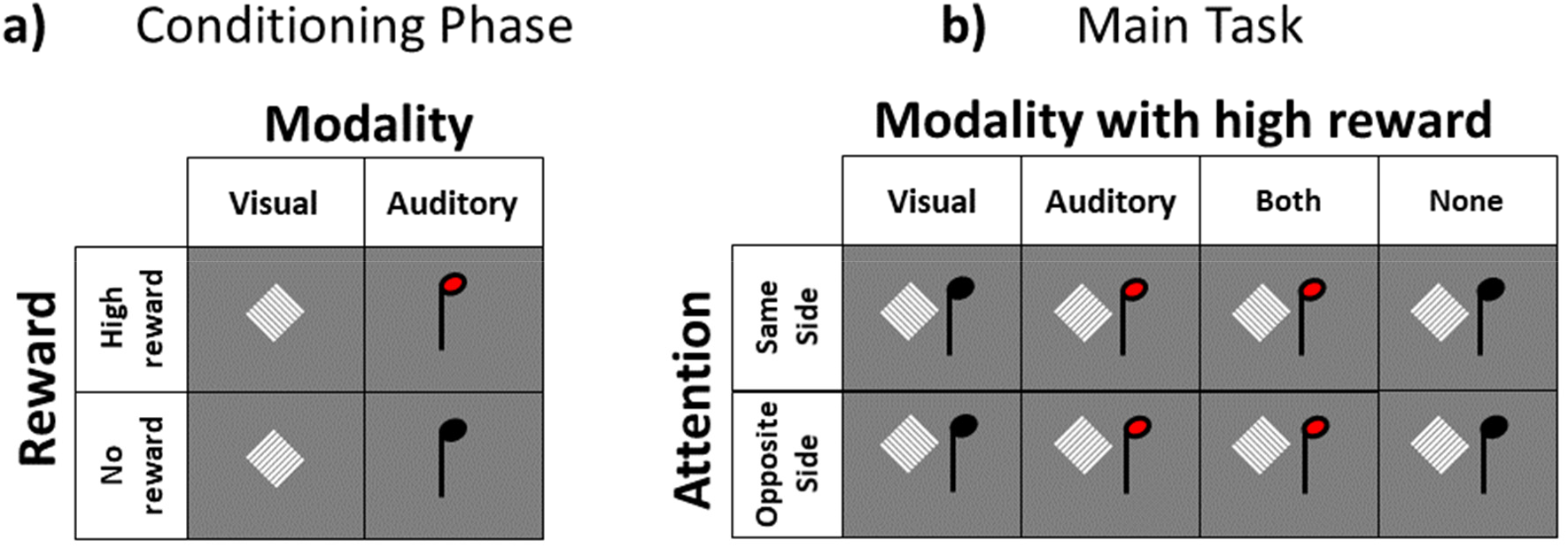
Stimuli and design of the experiment. **a)** Conditioning phase contained two independent factors: stimulus modality (visual or auditory) and reward value (high reward or no reward). **b)** In the main task two independent factors were manipulated: modality with high reward value (visual, auditory, both, none) and attention (same side or opposite side relative to the audiovisual stimulus).

### Main Task

At the beginning of each block (180 trials) of the task, participants were instructed to fixate their eyes on the screen center and pay attention to only one side (either left or right, counterbalanced across blocks, **Figure 1b**). Participants were required to only report targets (i.e., audiovisual stimuli containing a change in intensity) presented on the attended side.

All trials during the main task contained bimodal stimuli. We employed a 2 by 4 design yielding eight conditions which differed in the locus of attention (attended versus unattended) and reward value (reward on one modality, on both or on none). Specifically, the different reward value configurations consisted of: (1) visual high-value and auditory no-value: *VHSN* (2) visual no-value and auditory high-value: *VNSH* (3) visual high-value and auditory high-value: *VHSH* (4) visual no-value and auditory no-value *VNSN* (see **Figure 2b**). Participants completed 3200 trials during the main task where each of the reward conditions was repeated 400 times (200 trials per screen side). Of these, target trials (only on the instructed location) constituted 10% of all trials (i.e., 40 trials).

In both phases, a no-stimulus condition was added with the same number of repetitions as the other conditions (conditioning:240 trials and main task: 400 trials). In no-stimulus trials only the fixation point was displayed on the screen for the entire duration of the trial without any physical stimulus being presented. This condition was added to eliminate the overlapping electrophysiological activity elicited by the rapidly presented consecutive events (Woldorff, 1993), as implemented by (Talsma and Woldorff, 2005).

### Apparatus

The experiment was conducted in a darkened, sound-attenuated, and electromagnetically shielded chamber. Participants were seated on a chair with their heads fixed on a chinrest positioned 80 cm from a 22.5’ monitor (refresh rate = 120 Hz). Stimulus presentation was controlled by a PC under the Windows operating system equipped with MATLAB (version R2015b) and Psychophysics toolbox-3 (Brainard, 1997).

This study assessed the accuracy and reaction times (RT) of the detection of target stimuli (containing a transient change) based on participants responses indicated by a keypress on a keyboard. Eye movements were recorded with an EyeLink 1000 eye tracker system (SR Research, Ontario, Canada) in a desktop mount configuration, recording the right eye at a sampling rate of 1000 Hz. The electrophysiological (EEG) data was continuously recorded from 64 electrodes with an actiChamp system and referenced to the A2 electrode. The recording was done with BrainVision software (BrainVision Recorder 1.23.0001 Brain Products GmbH, Gilching, Germany; actiCap, Brain Products GmbH, Gilching, Germany). The signal was digitized at 1000 Hz and amplified with a gain of 10,000. All electrode/skin impedances were kept below 10kΩ.

### Transformations

#### Behavioral data

Reaction times (RTs) of correctly detected targets were computed separately for each condition at each phase of the experiment. Mean reaction time was used as the independent variable to assess the effect of reward and attention.

#### ERP analysis

EEG data was imported and processed offline using EEGLAB (Delorme and Makeig, 2004), an open-source toolbox running under the MATLAB environment. Raw data of each participant was band-pass filtered with 0.1 Hz as the high-pass cutoff and 40 Hz as the low-pass cutoff frequencies. An automatic bad channel detection algorithm was applied using EEGLAB’s pop_rejchan method (threshold = 5, method = kurtosis). All trials detected online as a keypress or fixation breaks (eye position to the fixation point > 9°) were removed from the analysis. After this, epochs were extracted using a stimulus-locked window of 3000 ms (1000 to 2000 ms). The remaining data were subjected to an Independent Component Analysis (ICA) algorithm (Delorme and Makeig, 2004). Eye-blinks and eye-movements artifacts were automatically identified and corrected using an ICA-based automatic method, implemented in the ADJUST plugin of EEGLAB (Mognon et al., 2011). Bad channels were interpolated by using the default spherical interpolation method. Data were re-referenced offline to the average reference. Finally, stimulus-locked epochs were extracted using a window of 1100 ms (-100 to 1000 ms), and baseline corrected using the pre-stimulus time interval (-100 to 0 ms). The stimulus-locked epochs were extracted using a window of 2200 ms (-1000 to 1200 ms), and baseline corrected using the pre-stimulus time interval (-100 to 0 ms).

Our ERP analysis was focused only on standard trials, i.e., trials *without* a change, as done previously (Talsma and Woldorff, 2005). Visual ERPs were measured in occipital electrodes (PO7/8 and O1/2) for P1 (70-170 ms) component. Auditory ERPs were measured at Fz for the auditory N1 (90-160 ms) component. P300 (250-400 ms) responses were measured over Pz. Analysis of visual P1 and auditory N1 components was based on the average within 60 ms window around the respective peak of each component measured across participants for each condition. Before the ERP analysis, we removed the overlapping ERP activity from the adjacent trials by subtracting a waveform of a no-stimulus condition from a waveform of each condition with an audiovisual cue, as Talsma et al. (2005) implemented.

Note that based on the visualization of the ERP traces, we changed the following measures compared to the pre-registered plan: firstly, we changed the auditory N1 window from 70-170 ms to 90-160 ms. Secondly, we increased the length of the time window around the peak of visual P1 and auditory N1 components within which ERP amplitudes were averaged from the pre-registered length of 30 ms to 60 ms. Thirdly, we changed the P300 window from 350-600 ms to an earlier time window 250-400 ms. Additionally, to be consistent with the method employed by (Talsma and Woldorff, 2005) visual ERPs were inspected in the contralateral electrodes, while we also report the results when ipsilateral visual ERPs were inspected (see the Supplementary information).

### Analysis

#### Behavioral data

Statistical analyses were done separately for the conditioning phase and the main task. In the conditioning phase, a two-way repeated-measures analysis of variance analysis (ANOVA) of RTs was done with factors *modality* (visual or auditory) and *value* (high-value cue or no-value cue).

In the main task (**Figure 3**), all stimuli were bimodal, where individual sounds and visual stimuli were either associated with high-value or no-value (VHSH – visual cue has high-value and sound has high-value; VHSN – visual cue has high-value and sound has no-value; VNSH – visual cue has no-value and sound has high-value; VNSN – visual cue has no-value and sound has no-value). Here a two-way ANOVA was used with the factor *auditory reward value* (high or no) and *visual reward value* (high or no). Significant main or interaction effects were subsequently inspected using planned pairwise comparisons.

**Figure 3.**
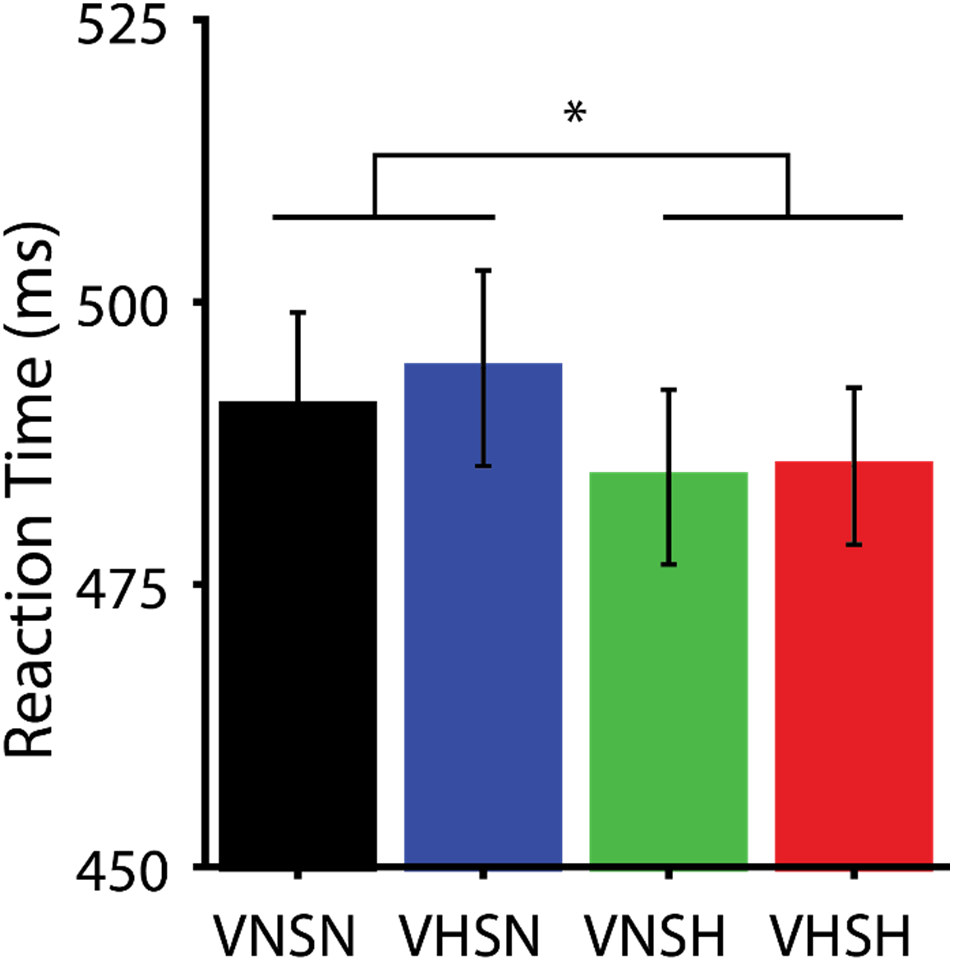
Reaction times during the main task. Four stimulus configurations consisted of conditions with high reward value in both modalities (VHSH), only in visual modality (VHSN), only in auditory modality (VNSH), or in neither of the modalities (VNSN).

#### EEG data during the main task

Firstly, to test whether we replicate the reported effects of a previous study that had inspired our paradigm (Talsma and Woldorff, 2005), ERP responses to the attended and unattended stimuli across all conditions were compared (**Figure 4**).

**Figure 4.**
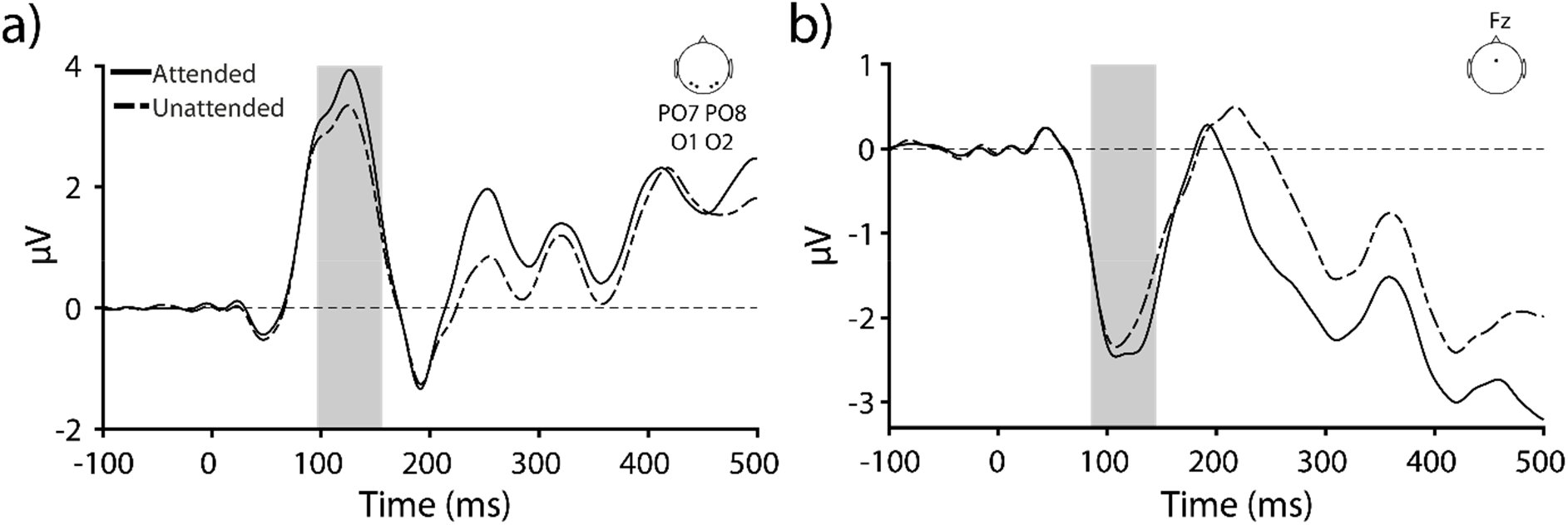
Attentional modulation of ERPs. **a)** ERP waves produced by the attended and unattended audiovisual stimuli that were recorded over contralateral occipital electrodes (PO7, O1, O2, PO8) or **b)** frontal electrodes (Fz).

Subsequently, a rmANOVA comprising factors *attention* (attended or unattended) and *value* (high- or no-value) was done on the data of conditions in which both visual and auditory stimuli of bimodal stimuli had the same value, i.e., either high- or no-value (VHSH and VNSN). Here, a main effect of reward would confirm *hypothesis 1*, a main effect of attention would confirm *hypothesis 2*, the absence of interaction effects would confirm *hypothesis 3* of our pre-registered plan (**Figure 5**).

**Figure 5.**
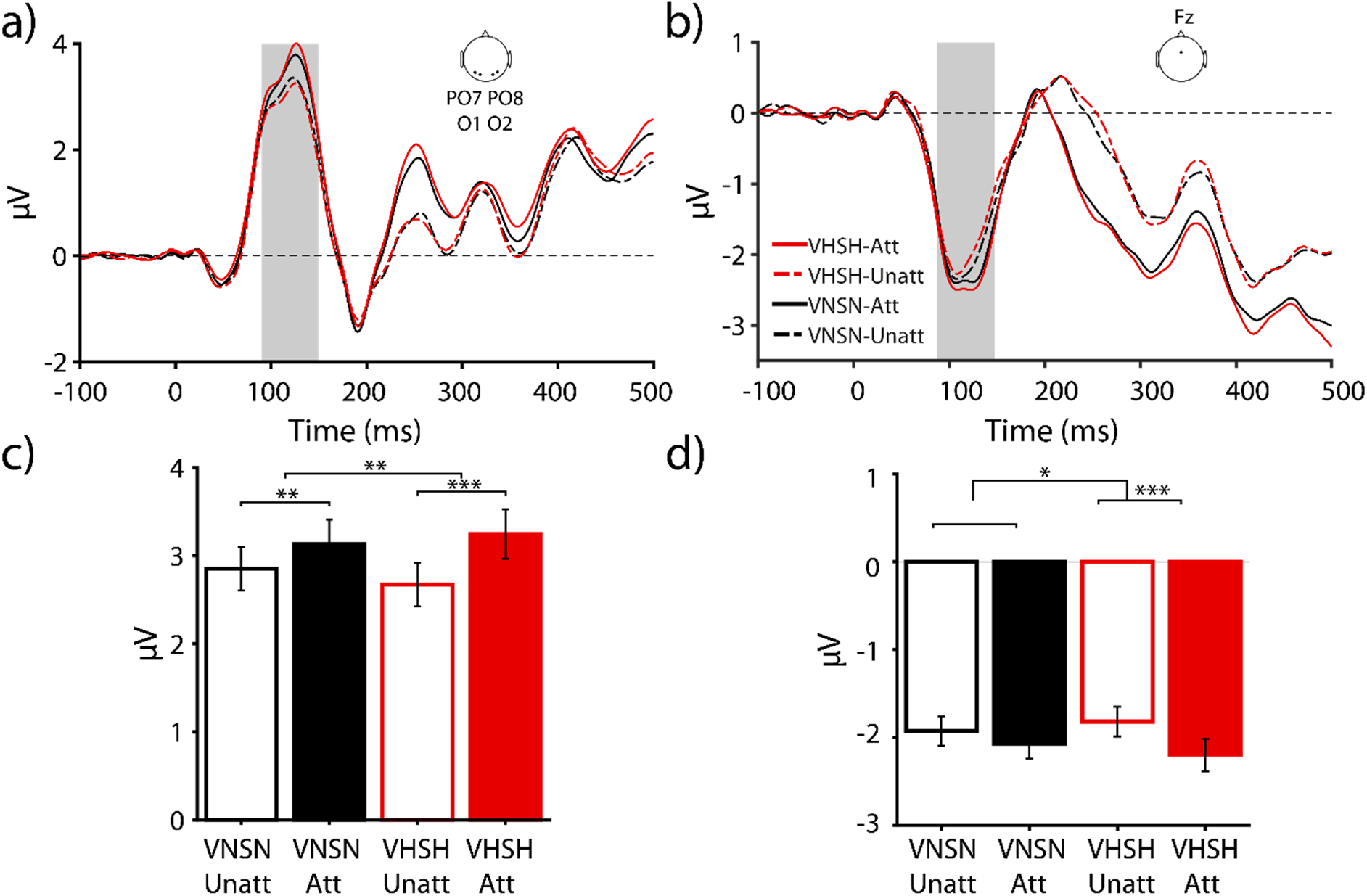
Modulation of early ERP components by attention and reward factors. ERP waves measured over contralateral occipital **a**) or frontal **b**) regions for conditions that either had high reward value in both modalities (VHSH) or no reward values in both modalities (VNSN) and were either presented on the attended (Att) or unattended (Unatt) side. **c)** Same as a) depicting the average visual P1 amplitudes. **d)** Same as b) depicting the auditory N1 amplitudes.

Having established an effect of attention in previous two analyses, we next subtracted the ERPs of attended and unattended conditions and tested the effect of reward value separately in each modality (**Figure 6**). To this end, a two-way rmANOVA with factors *value in visual modality* (high-value or no-value), *value in auditory modality* (high-value or no-value) was done separately on visual P1 and auditory N1 components (*hypothesis 4-6)*.

**Figure 6.**
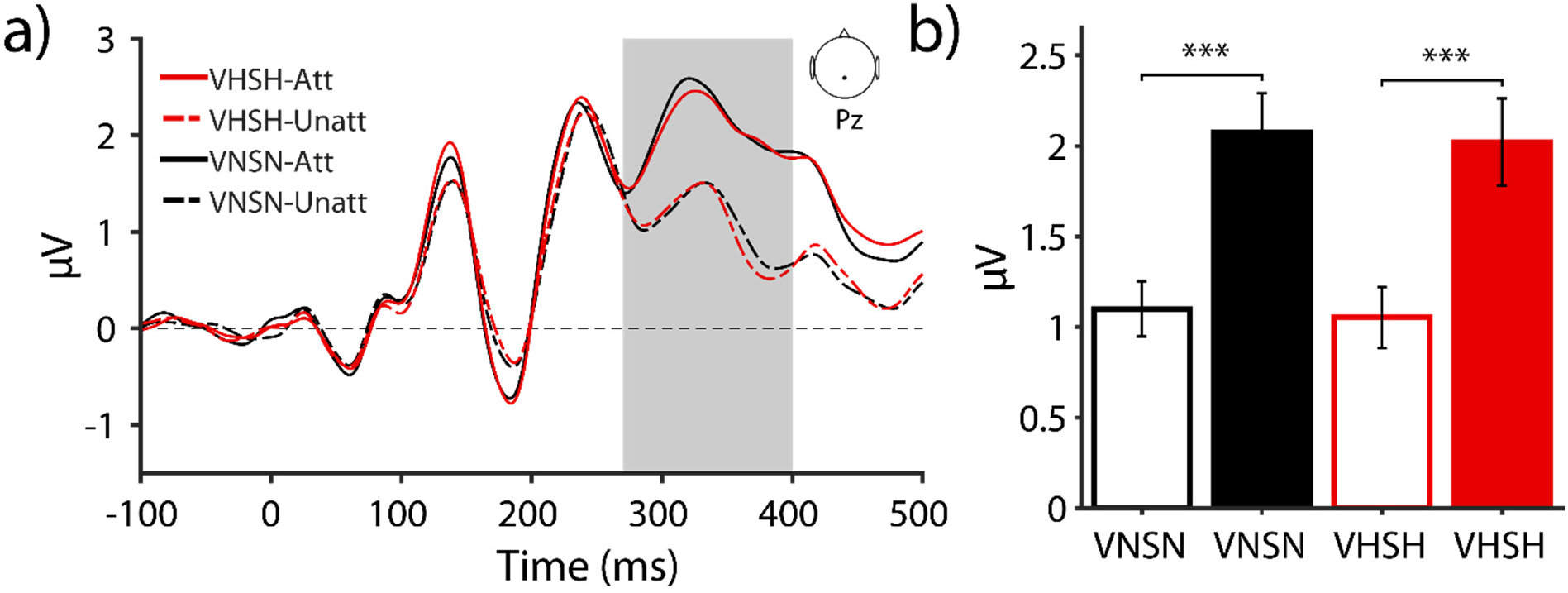
Modulation of late ERP components by attention and reward factors. **a**) ERP waves measured over Pz for conditions that either had high reward value in both modalities (VHSH) or no reward values in both modalities (VNSN) and were either presented on the attended (Att) or unattended (Unatt) side. **b**) average amplitude of P300 (250-400ms).

Finally, to test whether the effect of value in visual or auditory modalities differs between different regions of interest (ROI), we conducted a three-way rmANOVA with factors *ROI* (two levels: visual (PO7, O1, O2, PO8) or frontal region (Fz)), *value in visual modality* (high-value or no-value), *value in auditory modality* (high-value or no-value). Since P1 and N1 components have an opposite amplitude polarity, the amplitude of N1 component was rectified (multiplied by -1) before being included in this analysis. Here, our focus was on testing whether an i*nteraction* existed between value in each modality and the ROI (*hypothesis 7*).

## Results

### Behavioral results

The debriefing results and the analysis of behavioral and ERP responses (P300 component) during the conditioning indicated that participants had successfully learned the reward associations (see the Supplementary Information and **Figure S1**).

Behavioral results from the main task are shown in **Figure 3**. Participants had overall near perfect performance (Hit rate: mean ± s.e.m.: 94.72% ± 0.94) during the main task. Here, our main interest lay in how previously rewarded visual and auditory stimuli affected the speed of processing of the bimodal stimuli. Inspection of reaction times (see **Table 1**) revealed that conditions with a high-value auditory stimulus (VHSH and VNSH) had shorter reaction times compared to those with a high-value visual stimulus (VHSN and VNSN). A two-way rm ANOVA with the factor *visual reward value* (2 levels) and *auditory reward value* (2 levels) revealed only a main effects of *auditory reward value* F(1,37) = 5.36, *p* < 0.05, ηp2 = 0.127. Other main and interaction effects did not reach statistical significance (p>0.1).

**Table 1.**
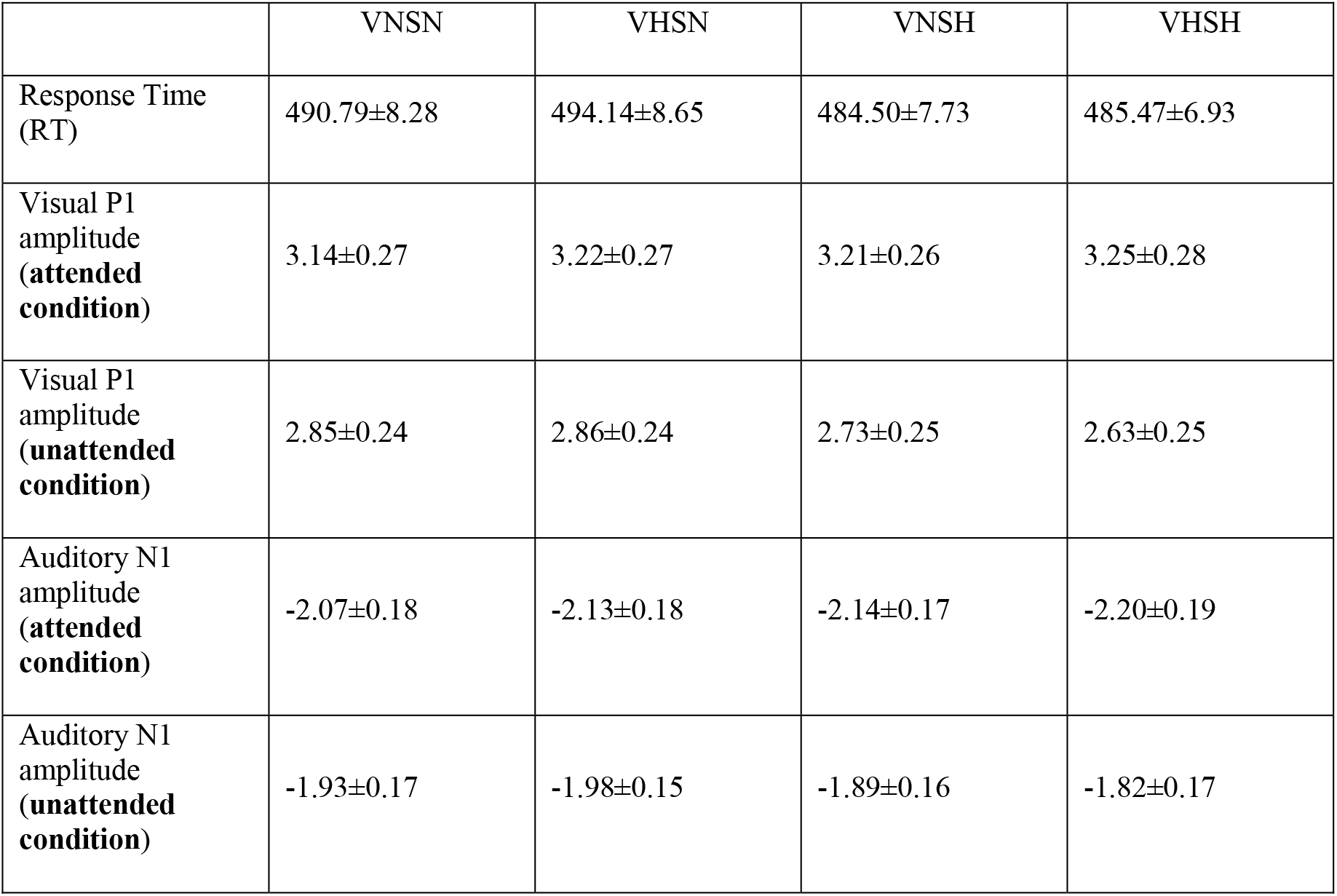
Overview of behavioral and electrophysiological indices measured during the main task in each stimulus condition.

|  | VNSN | VHSN | VNSH | VHSH |
| --- | --- | --- | --- | --- |
| Response Time (RT) | 490.79±8.28 | 494.14±8.65 | 484.50±7.73 | 485.47±6.93 |
| Visual P1 amplitude (**attended condition**) | 3.14±0.27 | 3.22±0.27 | 3.21±0.26 | 3.25±0.28 |
| Visual P1 amplitude (**unattended condition**) | 2.85±0.24 | 2.86±0.24 | 2.73±0.25 | 2.63±0.25 |
| Auditory N1 amplitude (**attended condition**) | -2.07±0.18 | -2.13±0.18 | -2.14±0.17 | -2.20±0.19 |
| Auditory N1 amplitude (**unattended condition**) | -1.93±0.17 | -1.98±0.15 | -1.89±0.16 | -1.82±0.17 |

Together, the analysis of behavioral data in conditioning and main task indicates a robust influence of reward value cued through the auditory modality on the target detection.

### Electrophysiological markers of attention and reward value

#### Attentional modulation of ERP responses

We first tested whether the attentional modulation effects reported by Talsma et al. (2005) are replicated in our paradigm, as shown in **Figure 4**. A one-way ANOVA on visual P1 component (PO7, O1, O2, PO8) revealed a main effect of *attention* (F(1,37) = 41.17, *p* < 0.001, ηp2 = 0.53). Attended conditions increased the amplitude of P1 component (mean ± SD: 3.20±0.27) compared to the unattended conditions (mean ± SD: 2.78±0.24). A similar effect was found in auditory N1 component measured at Fz (F(1,37) = 15.53, *p* < 0.001, ηp2 = 0.30), where attended stimuli increased the ERP amplitude of N1 component (mean ± SD: -2.13±0.17) compared to unattended stimuli (mean ± SD: -1.90±0.15). These results replicate previous findings (Talsma and Woldorff, 2005) and indicate that our experimental design modulated the allocation of spatial attention successfully.

### Interaction between attention and reward and modulation of ERP responses

#### Visual P1 component

Next, we examined the attentional and reward effects in conditions in which both modalities had either high- or no-value (VHSN and VNSN, see **Figure 5a,c).** Here, the analysis of visual P1 component revealed a main effect of attention (F(1,37) = 24.08, p < 0.001, ηp2 = 0.39) but no main effect of reward value (F(1,37) = 0.13, p = 0.71). Attended stimuli evoked higher amplitude of P1 component (mean ± s.e.m.: 3.19 ± 0.27), compared to unattended stimuli (mean ± s.e.m.: 2.76 ± 0.24). Interestingly, the difference between attended and unattended conditions was larger for high-value (mean ± s.e.m.: 0.58 ± 0.10, t(37) = 5.73, p < 0.001, dz = 0.93) compared to no-value (mean ± s.e.m.: 0.28 ± 0.10, t(37) = 2.79, p < 0.01, dz = 0.45) stimuli, leading to a significant interaction effect between attention and reward (F(1,37) = 8.38, p = 0.006, ηp2 = 0.18).

#### Auditory N1 component

Similar to the P1 component, an analysis of N1 component (**Figure 5b, d**) revealed a main effect of *attention* (F(1,37) = 9.2, *p* < 0.005, ηp2 = 0.2), no effect of *reward value* (F(1,37) = 0.01, *p* = 0.91), and an *interaction effect between attention and reward value* (F(1,37) = 6.10, *p* < 0.05, η ^2^ = 0.142). Attended stimuli evoked a more negative N1 peak (mean ± s.e.m.: -2.13 ± 0.18) compared to unattended stimuli (mean ± s.e.m.: -1.87±0.16). Additionally, high-value stimuli resulted in a larger modulation of N1 amplitude by attention (mean ± s.e.m.: -0.38 ± 0.09, t(37) = 4.1, p < 0.001, dz = 0.66) compared to no-value stimuli (mean ± s.e.m.: -0.14 ± 0.10, t(37) = 1.34, p = 0.18, dz = 0.21). Notably, the difference between attended and unattended no-value stimuli did not reach significance (*p* = 0.188).

#### P300 component

We next inspected the P300 component measured in Pz electrode (270-400 ms, see **Figure 6**). We found a strong main effect of attention (F(1,37) = 56.43, p < 0.001, ηp2 = 0.60) but no main effect of *reward value* (F(1,37) = 0.67, p = 0.41) or interaction between reward value and attention (F(1,37) = 0.00, p = 0.96).

Together, the results obtained in early visual and auditory ERP components (visual P1 and auditory N1) showed a main effect of attention and an interaction between attention and reward. A stronger attentional modulation for high-value stimuli demonstrates that stimulus value influences the allocation of attention.

### Dependence of the reward-driven modulation of attention on the sensory modality of reward stimuli

We have so far shown a strong effect of attention on early ERP components that is also influenced by the stimulus reward value. We next asked whether this effect depends on the specific configuration of stimuli; i.e., the modality through which reward value is cued (**Figure 7**). To this end, the ERPs of attended and unattended conditions were subtracted from each other and entered into an rmANOVA with factors *visual reward value* and *auditory reward value*.

**Figure 7.**
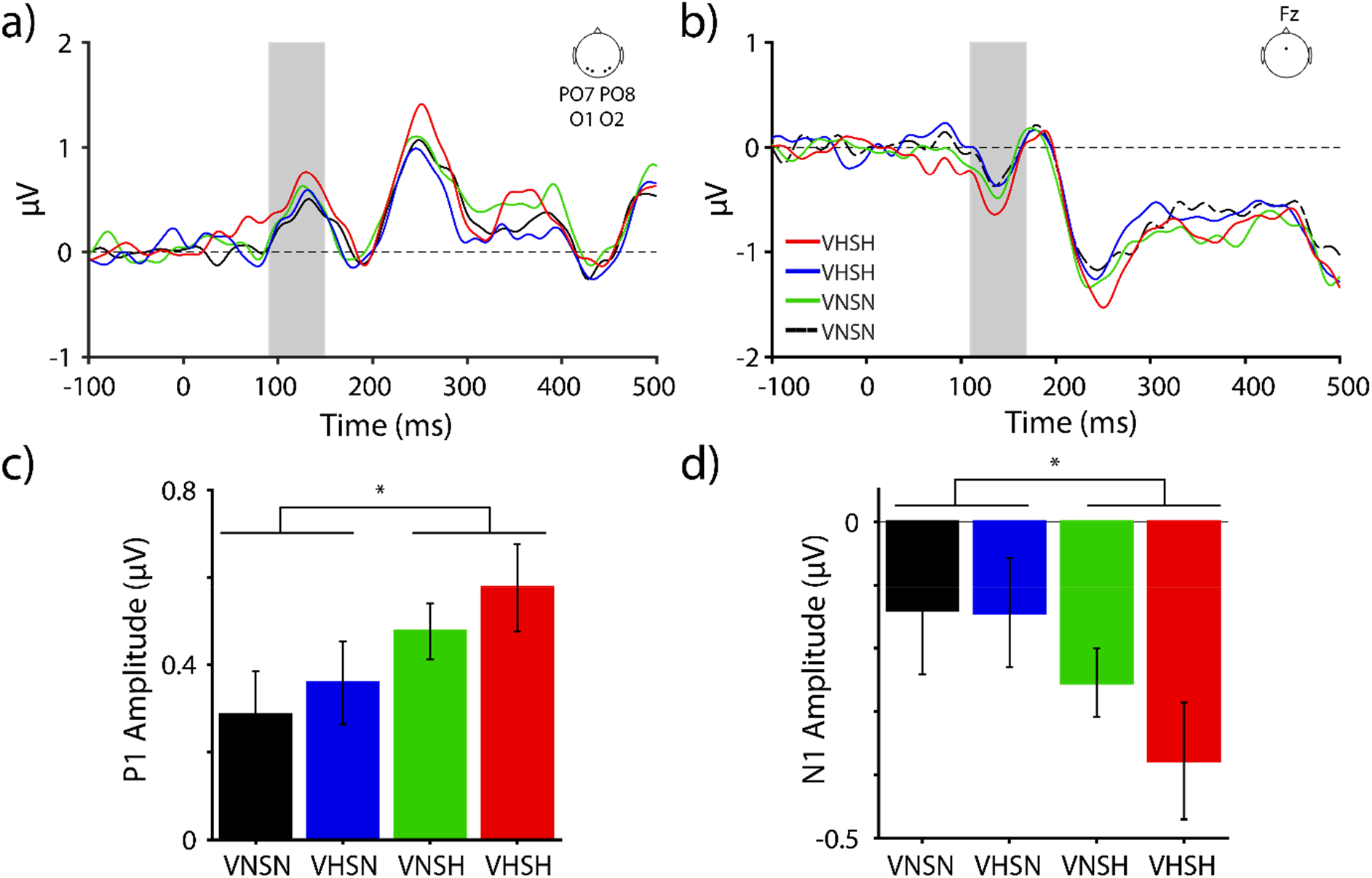
Modulation of attention by reward value of auditory and visual modalities. **a)** The difference wave between attended and unattended reward conditions (high reward in vision: VHSN, in auditory: VNSH, in both: VHSH, or in neither: VNSN modality) measured in contralateral occipital electrodes (PO7, O1, O2, PO8). **b)** Same as (**a**) for Fz electrode.

Analysis of visual P1 component revealed no main effect of *visual reward value* (F(1,37) = 1.61, *p* = 0.21, ηp2= 0.04) but a strong main effect of *auditory reward value* (F(1,37) = 9.74, *p* = 0.003, ηp2= 0.21). No *interaction* effect was found between these factors (F(1,37) = 0.02, *p* = 0.82). Importantly, the modulation of visual cortex by reward value was strongly lateralized and occurred predominantly in the contralateral sites with respect to the stimulus (see Supplementary **Figure S2**).

Analysis of auditory N1 component showed a main effect of *auditory reward value* (F(1,37) = 5.64, *p* = 0.023, ηp2= 0.13) as the presence of high-value auditory stimuli increased the N1 amplitude (mean ± s.e.m.: -0.32 ± 0.05 µV) compared to conditions that contained a no-value auditory cue (mean ± s.e.m.: -0.14 ± 0.08 µV). The main effect of *visual reward value* (F(1,37) = 0.94, *p* = 0.34) and the interaction of visual and auditory reward value were not significant (F(1,37) = 0.57, *p* = 0.45).

Inspection of individual stimulus configuration (see **Table 1** and **Figure 7**) reveals that both P1 and N1 had the highest amplitude when visual and auditory stimuli were associated with high-value (VHSH), followed by high-value only in auditory modality (VNSH) whereas other configurations (VHSN and VNSH) elicited lower amplitudes. Pairwise comparisons on visual P1 and auditory N1 component showed that VHSH elicited significantly higher amplitudes compared to VNSN (P1: Δ = 0.49, t(37) = 2.89, *p* = 0.006, dz = 0.47; N1: Δ = 0.24, t(37) = 2.47, *p* = 0.018, dz = 0.4) for both components and additionally was significantly higher than VHSN (Δ = 0.23, t(37) = 2.23, p = 0.032, dz = 0.36) for auditory N1. Other pairwise comparisons did not reach significance level (all *ps* > 0.1).

Finally, to test whether the influence of auditory and visual reward value depended on the region of interest (ROI), we conducted a rmANOVA on the visual P1 and auditory N1 components. This analysis revealed a main effect of *ROI* (F(1,37) = 10.06, *p* < 0.001, ηp2 = 0.21) as expected, and a main effect of *auditory reward value* (F(1,37) = 10.01, *p* < 0.005, ηp2 = 0.21). Importantly, no interaction was found between the auditory or visual reward value and ROI (F(1,37)<1, p>0.1), indicating that auditory reward value enhanced sensory processing both intra-modally (i.e., in Fz) as well as cross-modally (in occipital ROI).

## Discussion

The current study examined the interaction between spatial attention and reward value when visual, auditory, or both modalities were associated with high reward. During the conditioning phase, both visual and auditory stimuli sped up the target detection, with a stronger effect in auditory modality. During the main task, we found a strong effect of attention on early event-related responses of occipital and frontal areas (visual P1 and auditory N1 components, respectively), thus replicating a previous study that inspired our design (Talsma and Woldorff, 2005). We did not find a main effect of reward value, but importantly a significant interaction was found as attentional modulation of P1 and N1 responses were stronger when both modalities had high-value compared to no-value. These findings suggest that reward value modulates the strength of attentional filtering. However, when visual and auditory reward effects were compared separately, a more robust reward modulation of attention was found for the latter both behaviorally and electrophysiologically, thus reflecting the dominant role of auditory reward signals in the guidance of attention. Hence, in the context of our paradigm stimuli with high value, especially in auditory domain, guide the attentional resources towards the attended location and withdraw them from the unattended location.

A host of previous studies have demonstrated that reward biases selective attention towards stimuli previously associated with a positive outcome, an effect observed for shapes (Della Libera and Chelazzi, 2006, 2009), faces (Raymond and O’Brien, 2009), orientations (Kiss et al., 2009), and semantic information in Stroop tasks (Krebs et al., 2010). However, it is not always possible to determine the degree to which the observed behavioral or neural modulations are due to attention, reward, or both (Maunsell, 2004). Often reward-associated stimuli appear as the target of the task or contain target-related information, hence inadvertently engaging attention. More recent studies showed that even when reward stimuli are task-irrelevant (Garcia-Lazaro et al., 2018) or are delivered through a different sensory modality (Pooresmaeili et al., 2014; Antono et al., 2022) target processing is influenced by the reward information, suggesting independence of reward and attentional mechanisms. Specifically, a lack of influence of attentional load on reward effects (Baldassi and Simoncini, 2011) and temporally distinct ERP modulations for reward and attention (Bayer et al., 2017) supported the idea that selective attention and reward can influence perception independently. Here, we tried to resolve this ambiguity but while we had predicted an independence of attention and reward, we found evidence to the contrary. In contrast to previous studies that showed a clear separation between effects of reward and attention, our results show that reward effects on early perception occur exclusively through the modulation of attention. We note several major differences between the latter studies and ours. Firstly, Baldassi et al. (Baldassi and Simoncini, 2011) employed a weak manipulation of spatial attention, i.e. by performing a secondary task at fixation, compared to ours where attention was strictly controlled in space (to be on one or the other side). The form of attentional manipulation by Baldassi and Simoncini may have allowed a boost of reward effects through attention, despite the conclusion of the study (Baldassi and Simoncini, 2011). Secondly, in both Baldassi et al. (Baldassi and Simoncini, 2011) and Bayer et al. (Bayer et al., 2017) reward stimuli were predictive of the delivery of reward during the main task, whereas we tested stimuli that were previously associated with reward value and did not lead to rewards anymore. Reward predicting incentives can overall exert stronger effects on perception as better perceptual performance is constantly reinforced by the delivery of reward (Antono et al., 2022). This however does not allow to study the pure role of the *associated reward value* independent of any task-related feedback or continuous reward reinforcement. However, in the light of these studies and the fact that the reward effects on their own were weak in our paradigm, we surmise that independent effects of attention and reward can only be observed under conditions where reward can exert its maximal influence on perception without any assistance from attentional mechanisms. In other contexts, such as ours where rewards are learned prior to the task and are not further reinforced, the associated reward value of stimuli may affect perception primarily through the modulation of attention.

An important novel aspect of the current study is to investigate the effects of visual and auditory stimuli that were previously associated with high monetary reward outcomes. Previous studies on value-driven mechanisms have been mostly focused on reward effects in the visual modality (Failing and Theeuwes, 2018), although reward effects have also been reported in other sensory modalities (Rutkowski and Weinberger, 2005; Shuler and Bear, 2006; Pleger et al., 2008; Weil et al., 2010; Goltstein et al., 2013; Stanisor et al., 2013). This leaves a question of the extent to which value-driven mechanisms reflect a general principle of information processing across sensory modalities. A few recent studies approached this question by testing cross-modal reward modulations (Pooresmaeili et al., 2014; Anderson, 2016b; Sanz et al., 2018; Antono et al., 2022), testing the competition of reward and attention in other sensory modalities (Anderson et al., 2011a; Baines et al., 2011; Yantis et al., 2012; Chelazzi et al., 2014; Failing et al., 2015; Asutay and Västfjäll, 2016; Bourgeois et al., 2016; Tankelevitch et al., 2020; Kim et al., 2021; Qin et al., 2021), and comparing intra- and cross-modal reward modulation directly (Bruns et al., 2014; Pooresmaeili et al., 2014; Anderson, 2016b; Kang et al., 2017; Cheng et al., 2020; Bean et al., 2021; Vakhrushev et al., 2021; Antono et al., 2022). The emerging picture from these studies is that value-driven mechanisms reflect a general information processing principle that operates across modalities.

One important principle across sensory modalities is that when task-irrelevant features are associated with high reward they may potentially interfere with the processing of the target (Anderson et al., 2021). In line with this, recent studies (Vakhrushev et al., 2021) showed an early suppression of visual ERPs when the task-irrelevant reward cue was in the visual modality (i.e., intra-modally), but surprisingly late ERP components and behavioral sensitivity were enhanced when reward was cued through the auditory modality (Pooresmaeili et al., 2014; Vakhrushev et al., 2021). Hence, reward information exerted different effects on sensory processing dependent on whether it was cued intra- or cross-modally suggesting that modality-specific attentional resources might contribute to value-driven effects. Interestingly, although we used a different task and explicitly controlled the amount of spatial attention towards reward stimuli, our results are in line with the interpretation provided by (Vakhrushev et al., 2021). Notably, although visual high-value stimuli on their own (i.e., in VHSN configuration) led to no reward modulation of visual P1, the joint presence of high value in both modalities (i.e., VHSH) enhanced the ERPs, above and beyond both VNSN and VNSH conditions, indicating that congruent reward in both sensory modalities strengthens the reward-driven modulation of attention (**Figure 7**). These two findings reveal two important stages of reward-driven modulation of sensory processing. The *first stage* occurs locally within each sensory modality. At this stage, reward information competes with attention in order to gain priority in sensory processing (Anderson et al., 2011b, 2021; Failing and Theeuwes, 2018). As reward information conveyed through vision could potentially compete with the target detection task, for instance by withdrawing resources from a relevant feature (luminance) and allocating them to an irrelevant feature (line orientation), the putative local competition of reward and attention can lead to a reduction of reward effects. However, the local competition can be overturned by rewards in another sensory modality, suggesting that at a *second stage*, the reward information is integrated across different sensory modalities (Wise, 2004; Bruns et al., 2014; Cheng et al., 2020) resulting in an additive effect of reward of the two sensory modalities and a general improvement of sensory representations. Future studies will be needed to test the validity of this proposal.

We note that the lack of reward effects in visual modality could be due to two different reasons: (1) As mentioned above, these results may indicate a potential interference of reward information in visual modality with the allocation of spatial attention during the main task. Alternatively, (2) it is possible that participants had learned the associated reward value of auditory stimuli better, as also supported by the behavioral results obtained during the conditioning (see the Supplementary Information). Both scenarios are likely to have played a role and indicate that our effects may to some extent depend on the specific target detection task that we employed. Nevertheless, the strong effect of auditory reward on visual cortex remarkably corroborates the previous reports of cross-modal reward effects on perception.

Finally, in our experiment, participants learned the reward value of each sensory modality separately and they were then exposed to bimodal stimuli containing the same or different reward values in two modalities. The pattern of results observed here may be due to this training protocol, as auditory and visual stimuli were never paired during the learning. Alternatively, learning the associated reward value of a bimodal stimulus may promote integrating sensory features and reward values more strongly across sensory modalities and thereby lead to reward modulations that have less dependence on modality-specific mechanisms. Future studies will be needed to examine the role of learning protocols in how reward signals from different sensory modalities interact.

## Conclusion

In summary, we found that the allocation of spatial attention towards audiovisual stimuli is guided by the associated reward value of auditory and visual modalities. In the context of the task employed here, value-driven modulation of attention was more robust in auditory modality, with the maximum effect observed when both visual and auditory components of an audiovisual stimulus were associated with high value. These results inspire a two-stage model in which reward information is first represented separately in each sensory modality and is subsequently integrated across modalities. The integration of reward value boosts the combined value of a bimodal stimulus and at the same time enhances the attentional selection of the task-relevant information.

## Acknowledgments

We thank Franziska Ehbrecht for her help with the data collection. This work was supported by an ERC Starting Grant (no: 716846) to AP.

## Supplementary Information

### Behavioral and Electrophysiological indices of reward learning

The direct assessment of reward associations during and after the conditioning phase (debriefing) indicated that participants successfully learned the reward values of both visual and auditory stimuli: all participants indicated correctly which of the two stimuli presented to them had been associated with a high value, and this was the case in both modalities.

Analysis of reaction times revealed a main effect of *modality* F(1,37) = 13.9, *p* < 0.001, ηp2 = 0.27 and a main effect of *value* F(1,37) = 12, *p* < 0.001, ηp2 = 0.25, as well as an interaction effect F(1,37) = 5, *p* = 0.031, ηp2 = 0.12. Participants were faster at detecting the visual targets (mean ± s.e.m.: 486.06 ± 6.49) compared to the auditory targets (mean ± s.e.m.: 506.41 ± 6.08). Similarly, participants were faster at detecting the change in high-value stimuli (mean ± s.e.m.: 490.97 ± 5.94) compared to the change in no-value stimuli (mean ± s.e.m.: 501.49 ± 5.79). Follow-up pairwise comparisons revealed that the reward effects were strongest for the auditory stimuli (t(37) = -3.25, *p* < 0.005, dz = 0.53), where response to a high-value auditory cue (mean ± s.e.m.: 497.25 ± 6.40) was much faster than to a no-value auditory cue (mean ± s.e.m.: 515.56 ± 6.99). The difference between high- and no-value visual stimuli did not reach significance (t(38) = -0.83, *p* = 0.41, dz = 0.13).

Additionally, we inspected the P300 responses to assess the learning of reward associations (**Supplementary Figure S1**). This analysis again confirmed that participants had learned the reward associations, although the P300 modulation by reward was only significant for the auditory stimuli (main effect of reward value: F(1,37) = 53.09, *p* < 0.001, ηp2 = 0.59; interaction of reward and modality: F(1,37) = 41.54, *p* < 0.001, ηp2 = 0.53). Together, these results suggest that all participants had learned the reward associations of both modalities in our task, although the auditory rewards seemed to predominate in their strength of behavioral and electrophysiological modulations.

**Figure S1.**
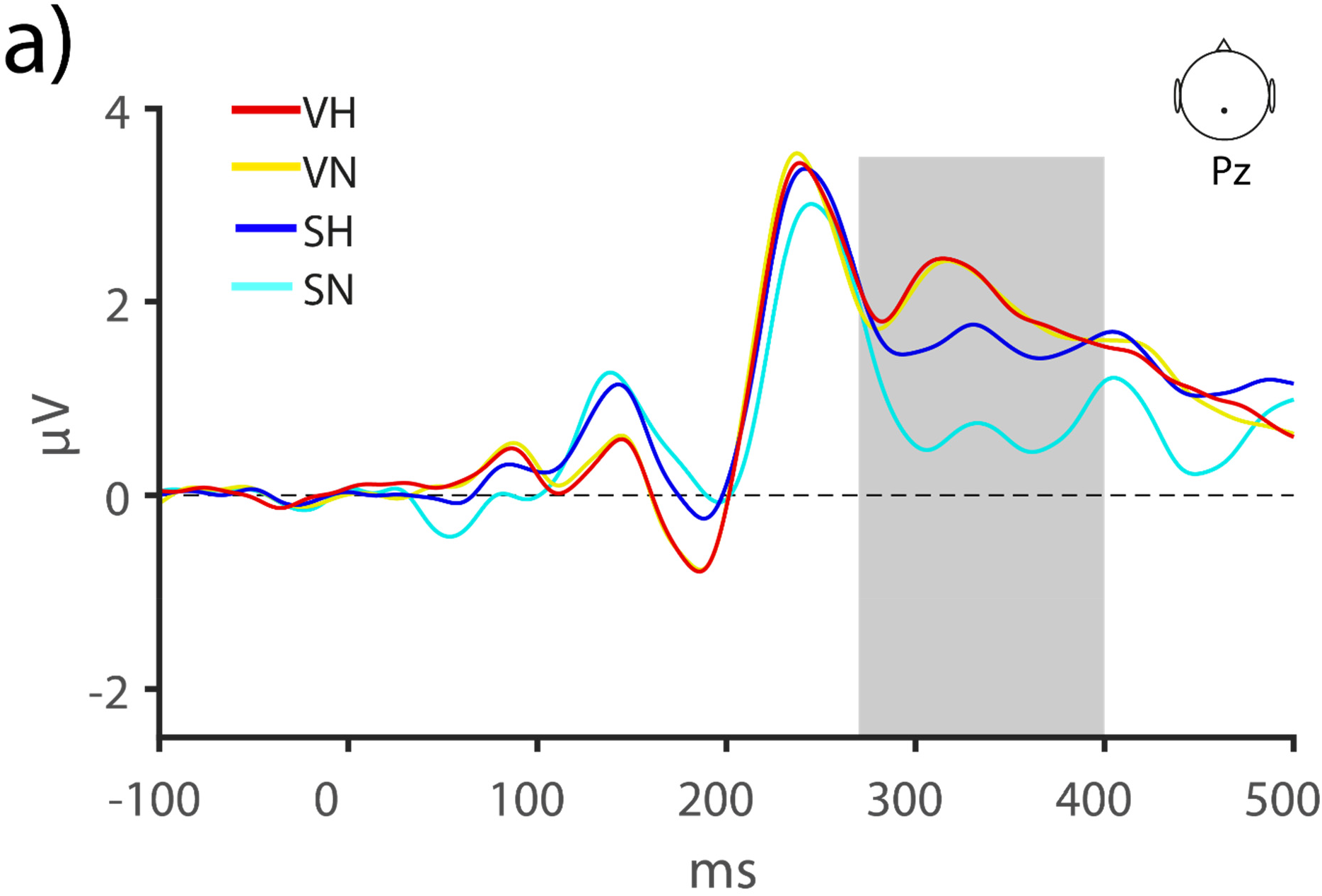
Reward modulation of P300 components by reward during the conditioning. ERP waves were measured over Pz for conditions with high value (VH or SH) or no reward values (VN or SN) and were tested for the average amplitude of P300 (270-400 ms).

### Reward-driven modulation of visual ERPs occurred primarily in contralateral sites

Due to the retinotopic organization of visual cortex, neuronal responses in each hemisphere are most strongly evoked by the stimuli that are presented in the contralateral visual hemifield. To test the dependence of the reward-driven effects on the laterality of the stimuli, we inspected the contra- and ipsi-lateral ERPs separately (**Figure S2**). An ANOVA revealed a trend for a significant interaction effect between *auditory reward value (high- or no-value)* and *hemisphere (contra- or ipsi-lateral)* F(1,37) = 3.06, *p* = 0.08, ηp2 = 0. 076. The reward-driven modulation of attention was strongest in contralateral (Δ = 0.21 ± 0.06, *p* = 0.003) compared to ipsilateral (Δ = 0.06 ± 0.09, *p* = 0.49) ERPs.

**Figure S2.**
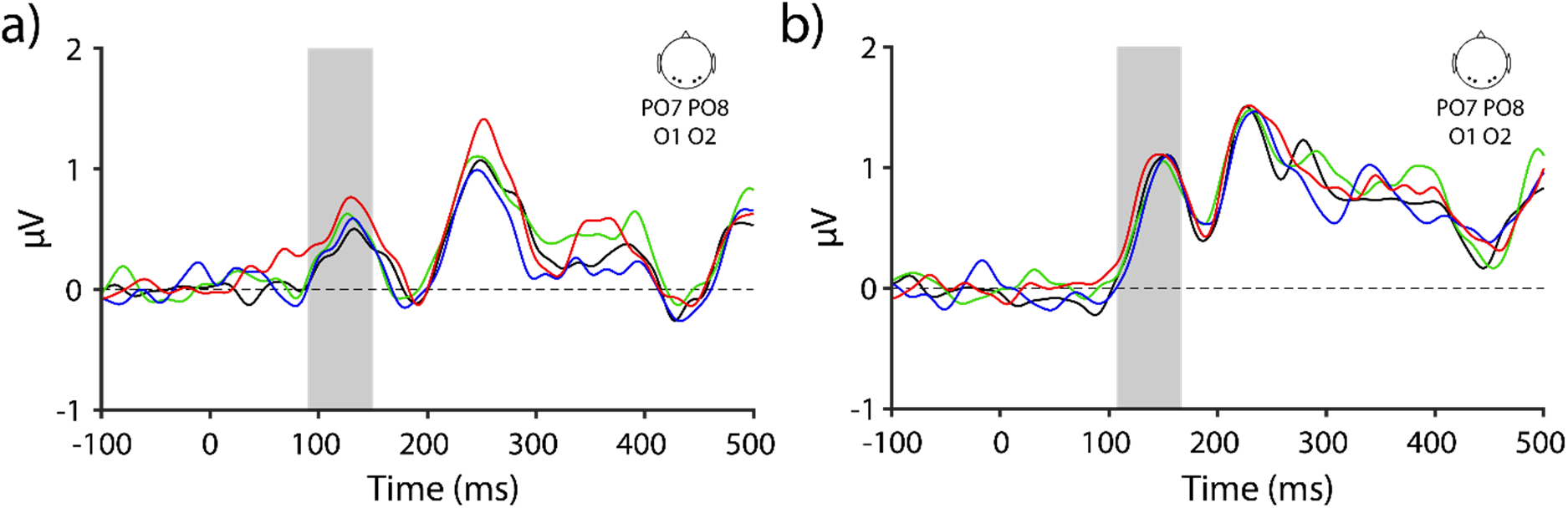
Modulation of attention by reward value of auditory and visual modalities in ipsilateral and contralateral visual cortex. **a**) The difference waves between attended and unattended reward conditions (high reward in vision: VHSN, in auditory: VNSH, in both: VHSH, or in neither: VNSN modality) measured in contralateral occipital electrodes (PO7, O1, O2, PO8). **b**) Same as (**a**) for ipsilateral occipital electrodes (PO7, O1, O2, PO8)

